# Exploring the Flexible Penalization of Bayesian Survival Analysis Using Beta Process Prior for Baseline Hazard

**DOI:** 10.1101/2024.09.20.614120

**Authors:** Kazeem A. Dauda, Ebenezer J. Adeniyi, Rasheed K. Lamidi, Olalekan T. Wahab

**Affiliations:** Department of Mathematics, University of Bergen, 5007 Bergen, Norway; Department of Mathematics and Statistics, Kwara State University, Malete, Nigeria; alt-Address: Department of Mathematics and Statistics, Kwara State University, Malete, Nigeria

**Author notes:** Corresponding author, (Kazeem A. Dauda). Email addresses:* (Ebenezer J. Adeniyi), (Rasheed K. Lamidi), (Olalekan T. Wahab).

**Keywords:** penalization function, semiparametric model, shrinkage parameter, joint posterior function, MCMC Algorithm

## Abstract

High-dimensional data has significantly captured the interest of many researchers, particularly in the context of variable selection. However, when dealing with time-to-event data in survival analysis, where censoring is a key consideration, progress in addressing this complex problem has remained somewhat limited. Moreover, in microarray research, it is common to identify groupings of genes involved in the same biological pathways. These gene groupings frequently collaborate and operate as a unified entity. Therefore, this study is motivated to adopt the idea of a Penalized semi-parametric Bayesian Cox (PSBC) model through elastic-net and group lasso penalty functions (PSBC-EN-G and PSBC-GL-G) to incorporate the grouping structure of the covariates (genes) and optimally perform variable selection. The proposed methods assign a beta prior process to the cumulative baseline hazard function (PSBC-EN-B and PSBC-GL-B), instead of the gamma prior process used in existing methods (PSBC-EN-G and PSBC-GL-G). Three real-life datasets and simulation scenarios were considered to compare and validate the efficiency of the modified methods with existing techniques, using Bayesian Information Criteria (BIC). The results of the simulated studies provided empirical evidence that the proposed methods performed better than the existing methods across a wide range of data scenarios. Similarly, the results of the real-life study showed that the proposed methods revealed a substantial improvement over the existing techniques in terms of feature selection and grouping behavior.

## 1. Introduction

Survival analysis is dedicated to investigating the time until a specific event, commonly termed as a “failure” [1]. This scenario finds widespread application across diverse scientific domains, including medicine [2, 3], biology [4, 3], engineering [5], and economics [6, 7]. Researchers employ survival analysis to scrutinize the time leading up to events such as mortality or equipment malfunction [8]. For instance, researchers may investigate the survival times of cancer patients from the time of diagnosis, where the event of interest could be the occurrence of relapse or, unfortunately, the death of the patient[3, 8, 9, 10].

Recently, there has been increasing interest in modeling survival data using deep learning methods in medical research [11, 12, 13]. However, these deep neural network-based survival models provide only point estimates of the hazard rates and thus cannot properly convey uncertainty in the estimations. Hence, to properly consider the uncertainties in deep neural network-basd survival models. Several researchers have proposed a Bayesian deep learning approach [5, 14]. As an illustration, Feng, and Zhao [14] proposed a Bayesian hierarchical deep neural networks model that can provide not only point estimate of survival probability but also quantification of the corresponding uncertainty. In addition, [5] handle-censored, and capture the non-linear relationship between covariates and target variable lifetime.

At the same time, Bayesian approaches to variable selection have also become popular, not least since the relevance of a covariate can be assessed simply by computing the posterior probability that it is included in a model [15, 16]. This recast variable selection as a model selection problem with every possible model assigned an individual posterior probability. One of the most popular of such model selection priors is the spike-and-slab prior [2, 17]. Among of the advantages of the method is the ability to adjust concentration points for flexible prior distributions, resulting in nonlinear shrinkage of regression coefficients and smoothed estimates. In addition, it enables fully Bayesian inference through Markov chain Monte Carlo, and aims to create a unified framework for variable selection, producing sparse coefficient structures in both low and high-dimensional contexts [18, 19, 20, 21].

Interestingly, among of the several traditional modeling methods for survival analysis well-documented in the literature, the Cox proportional hazard (Cox-PH) model stands out as a prominent semiparametric technique [22, 23]. And, recognized for its practical applicability, it facilitates a flexible modeling approach with minimal assumptions by maximizing the partial likelihood and bypassing the need to model the baseline hazard function. Although, the Bayesian paradigm mandates an explicit parametrization of the baseline hazard function. However, extensive research has demonstrated the adoption of Gamma process [24, 25, 26], a prevalent non-parametric process prior, in Bayesian proportional hazards models to characterize the cumulative baseline hazard.

Similarly, Ibrahim et al.[25] proposed a semi-parametric approach, combining a non-parametric prior for the baseline hazard rate (using, a discrete gamma process prior for the baseline hazard) with a fully parametric prior for the regression coefficients. Notably, the method demonstrated effective performance with low-dimensional data [25, 27, 28, 29]. However, the methodology has yet to be applied to high-dimensional data, and its suitability for grouped data has not been explored.

Subsequently, while there have been studies on variable selection methods [30, 31, 32, 33, 34] pioneered a variable selection approach incorporating the widely recognized lasso penalty within the Cox proportional hazards (Cox-PH) model framework, where the likelihood function is grounded, and the cumulative baseline hazard function is modeled using a gamma process. The study introduced a prior on the tuning parameter for shrinkage, offering adaptive control over the model’s sparsity. Similarly, Lee et al.[35] applied a data augmentation approach to handle censored survival times and enhance prior-posterior conjugacy. Also, for identifying relevant grouped covariates, a shrinkage prior distribution for regression coefficients, emulating the effect of a group lasso penalty was assigned. And, lastly to address the challenge of not shrinking coefficient estimates to exact zeros in a Bayesian penalized regression approach, a two-stage thresholding method utilizing the scaled neighborhood criterion and Bayesian information criterion was employed. However, the method was found to outperformed the competing methods on both simulated and real-life data, in terms of variable selection accuracy and predictive power.

Hence, in this paper, considering the typical distribution patterns of survival analysis, which often manifest as either gamma or beta distributions, researchers have a reasonable basis to explore the incorporation of a beta process. Thus, there is a need to implement the beta perspective within the framework, representing a modification of the methodologies proposed by [36] and [37]. Hence, this study implements and adopts the beta prior for the baseline hazard for effective prediction and variable selection in high-dimensional survival data. A similar version of the beta prior was introduced in the master’s thesis of one of the authors of this study in 2023 [38]. In this phase, we present the mathematical formulation of the beta prior and the joint posterior for both elastic and group lasso, as well as real-life applications, respectively.

This paper is structured as follows: In Section 2.1, we present a comprehensive overview of the penalized semi-parametric Bayesian Cox model, incorporating grouping and shrinkage priors. Section 2.2.3 outlines the joint posterior distribution and the turning parameter in detail. Formulation of the modified joint posteriors is discussed in Section 2.3, while Section 2.4 provides a thorough computational algorithm for obtaining both the existing and proposed penalized semi-parametric Bayesian Cox model (PSBC). To assess the performance of our proposed methods, Section 3.1 present the simulation of the study. In Section 3.2, we apply the proposed models (PSBC-EN-B and PSBC-GL-B) alongside the existing models to analyze three distinct microarray gene expression datasets. Concluding this study, Section 4 presents our closing remarks on the study.

## 2. Methodology

In this section, we briefly review the general concept of the penalized semiparametric Bayesian Cox model, examining existing methods and introducing new methods that focus on the prior of the cumulative baseline hazard function, joint posterior distributions, and the full Bayesian technique for selecting tuning parameters. Moreover, we discuss the formulation of the sequential BIC as a thresholding method for both existing and proposed approaches.

### 2.1. Penalized Semiparametric Bayesian Cox Model with Grouping and Shrinkage Priors

Suppose that a data set consists of *n* subjects, and for the individual *i*^*th*^ subject we record the actual survival time *T*_*i*_, covariates *X*_*i*_ = (*X*_*i*1_, …, *X*_*ip*_)^*′*^ and the event indicator *δ*_*i*_ ∈ {0, 1}. In right censoring, 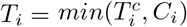 where 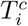 *and C*_*i*_ are survival and censoring times, respectively (*i* = 1, …, *n*), and nonnegative random variables; and the event indicator 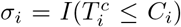 where *I*(.) is an indicator function. The data structure for this scenario was properly illustrated in table 1.

**Table 1:**
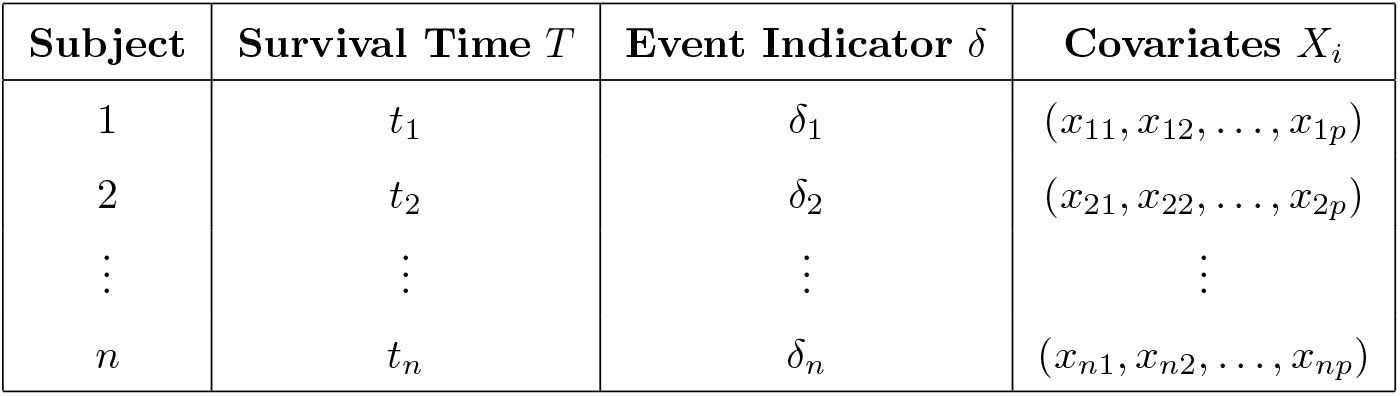
Survival times, event indicators, and covariates for all subjects.

By assuming that subjects are independent from each other and that 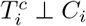 given *X*_*i*_. Thus, the conditional hazard function given *X*_*i*_ (*h*(*t/X*_*i*_)) quantifies the instantaneous failure rate at a given time *t* with *X*_*i*_. Hence, the model that links the conditional hazard function to *X*_*i*_ is denoted by equation (1), commonly called the Cox model [22], a well-known semiparametric model in survival analysis.

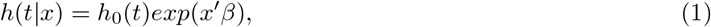

where *β* = (*β*_1_, …, *β*_*p*_)^*′*^ is a column vector of p regression parameters, and *h*_0_(*t*) is unspecified arbitrary baseline hazard function. Under the Cox model (1) the joint survival probability of *n* subjects given the matrix of covariates *X* is given by

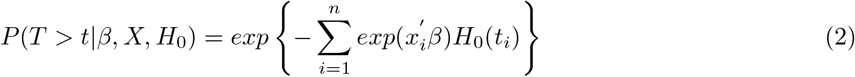

where *H*_0_(*t*_*i*_) is the cumulative baseline hazard function. A gamma process prior is assigned to the cumulative baseline hazard function *H*_0_(*t*)

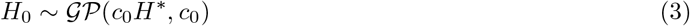

*H*^*\**^(*t*) is an increasing function with *H*^*\**^(0) = 0 and *c*_0_ is a positive constant. Let *h*_*j*_ denote the increment in the cumulative baseline hazard in the interval *I*_*j*_, as follows

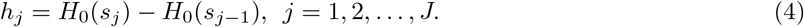

The gamma process prior in (4) implies that the *h*_*j*_ follows independent gamma distribution, that is

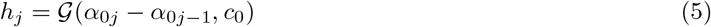

where *α*_0*j*_ = *c*_0_*H*^*\**^(*s*_*j*_). Therefore, the conditional probability of the *i*^*th*^ subject failing in the interval *I*_*j*_ is given by

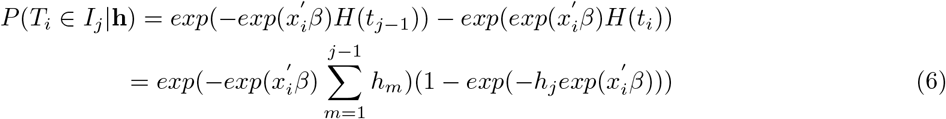

where *h* = (*h*_1_, *h*_2_, …, *h*_*j*_)^*′*^ This leads to our grouped data likelihood function

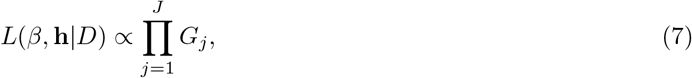

where 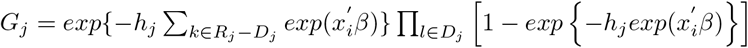,

The regression coefficients *β* = (*β*_1_, …, *β*_*p*_) plays a major role in selecting the covariates in the model (7) So we introduce a prior that will results in shrinkage in our model. The variable selection was done through BIC thresholding. Note that, the |*β*_*j*_| in lasso penalty is proportional to the (minus) log-density of Laplace distribution. Laplace prior (8) for the regression coefficients is given by,

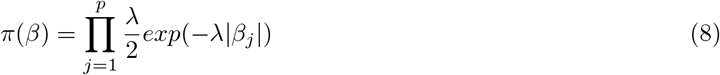

Meanwhile, we use the conditional Laplace prior in (9), instead of (8) in order to guarantee unimodality.

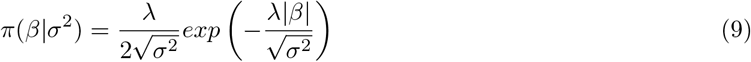

A noninformative marginal prior 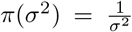 is assigned on *σ*^2^. The hierarchical representation of the Bayesian lasso with prior (9) can thus be obtained by utilizing the representation of the Laplace distribution as a scale mixture of normals with an exponential mixing density as described in (10).

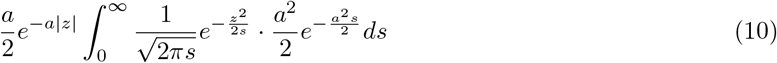

### 2.2. Overview of existing methods and parameter tuning

#### 2.2.1. PSBC Elastic Net (PSBC-EN-G)

The conditional *prior* of *β*|*σ*^2^ for the elastic net penalty can be expressed as

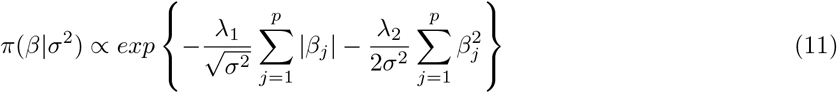

Combining (5), (7), (10), and (11), the hierarchical representation of our penalized semiparametric Bayesian Cox model with the elastic net prior (`PSBC-EN-G`) can be described as

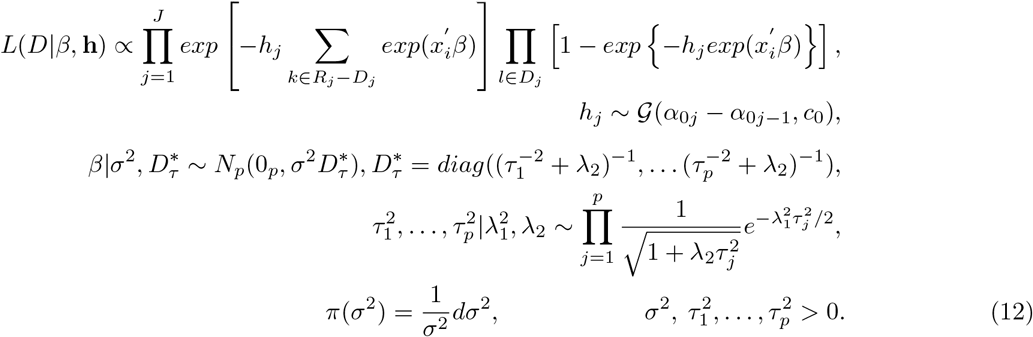

Note that the conditional priors of 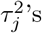 in (12) are not independent gamma distributions, 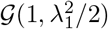 as given by [39]. The prior for 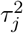 given in equation (12) is

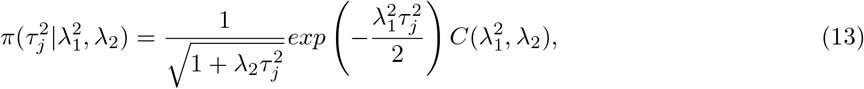

where

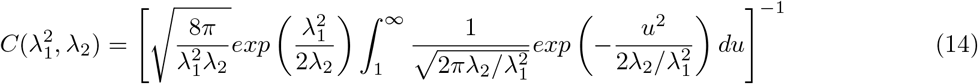

is the normalizing constant.

#### 2.2.2. PSBC Group Lasso (PSBC-GL-G)

The conditional prior of *β*|*σ*^2^ for the group lasso penalty can be expressed as

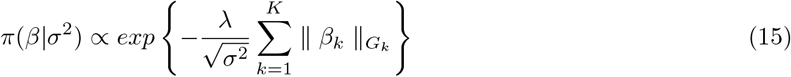

It is noted that the group lasso prior (15) assumes that variables are arranged in *K* groups, each of size *m*_*k*_. Integrating (5), (7), (10), along with the hierarchical formulation of our penalized semiparametric Bayesian Cox model and the fused lasso prior (PSBC-FL), the hierarchical representation of our penalized semiparametric Bayesian Cox model with the group lasso prior (PSBC-GL-G) can be described as

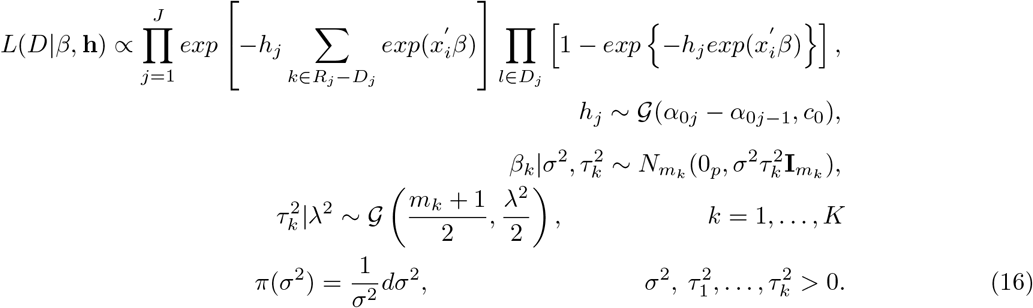

#### 2.2.3. Joint Posterior Distributions & Tuning Parameters

The tuning parameters in (12) and (16) that regulate the sparsity and grouping effect in the data set are *λ, λ*_1_, and *λ*_2_. The tuning parameters were selected using the full Bayes technique. Setting *λ*^2^ (in PSBC-GL-G) and 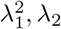(in PSBC-EN-G) a gamma prior in (17).

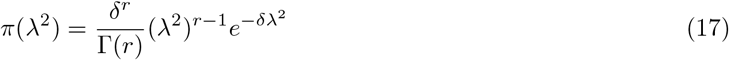

where *r*, and *δ* are the shape and the rate parameters of the gamma distribution. The prior distributions of tuning parameters are listed as

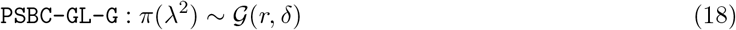

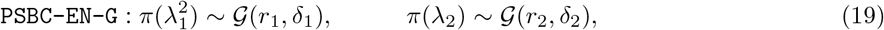

The model’s sparsity is determined by the *λ* (in `PSBC-GL`) and the *λ*_1_ (in `PSBC-EN-G`). Thus, if *λ* (or *λ*_1_) *→* 0, then all the covariates are chosen, and if *λ* (or *λ*_1_) *→ ∞* in `PSBC-FL`, then no covariates are included in the model. Adjacent parameters shrink toward the same value as a result of a more very strict constraint provided by the sequential differences of the regression parameters.

The final joint posterior distribution `PSBC-EN-G` model can be expressed as follows by combining (12), (13), (14), and (19).

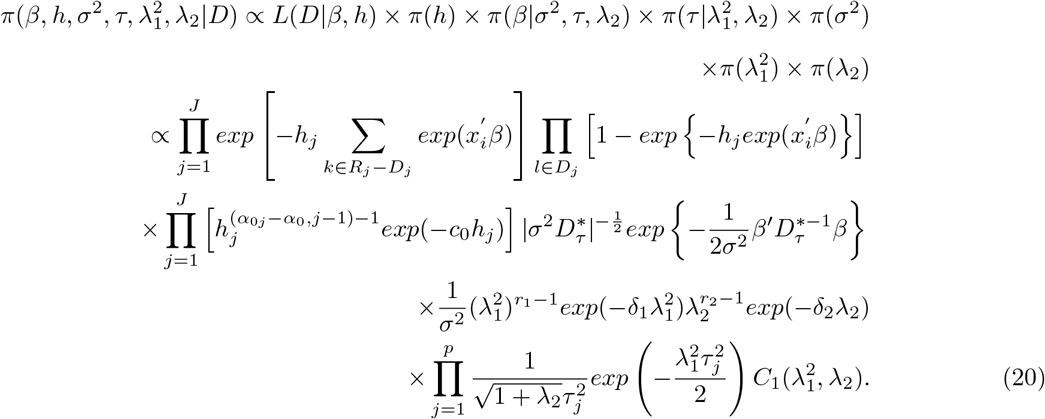

And, the final joint posterior distribution of our `PSBC-GL-G` model can be expressed as follows by combining (16) and (18).

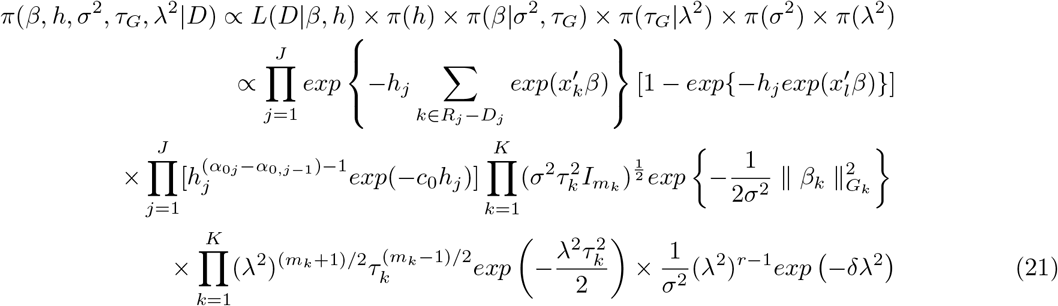

where *τ*_*G*_ = (*τ*_1_, …, *τ*_*k*_)^*′*^.

### 2.3. Formulation of proposed methods:PSBC-EN-B and PSBC-GL-B Joint Posteriors

This section presents the formulation of the joint posterior of PSBC-EN-B and PSBC-GL-B models. These models are constructed upon the grouped data likelihood (22), wherein the cumulative baseline hazard function in the Cox model is a priori modeled by a beta process. Moreover, the joint posterior of each model was based on full Bayesian approach for selecting the tuning parameters.

However, these posteriors aim to improve the accuracy and efficiency of parameter estimation in high-dimensional statistical modeling, especially in scenarios characterized by complex data structures and a plethora of predictors. Additionally, through the incorporation of grouping and shrinkage priors, this modification seeks to strike a balance between variable selection, coefficient estimation, and computational tractability.

#### 2.3.1. Development of PSBC Elastic Net (PSBC-EN-B) joint posterior

Assigning beta prior process to the cummulative baseline hazard function *H*_0_(*t*). The likelihood can be written as

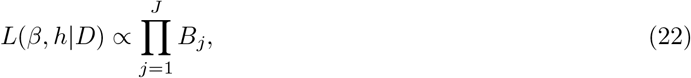

where 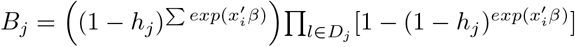.

where *h* = (*h*_1_, *h*_2_, …, *h*_*j*_)^*′*^. To complete the discretized beta process model, we specify independent beta priors for the *h*_*j*_’s. Specifically, we take

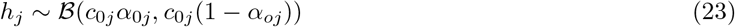

and independent for *j* = 1, 2, …, *J*.

Hence, the final joint posterior distribution of our `PSBC-EN-B` model can be written as,

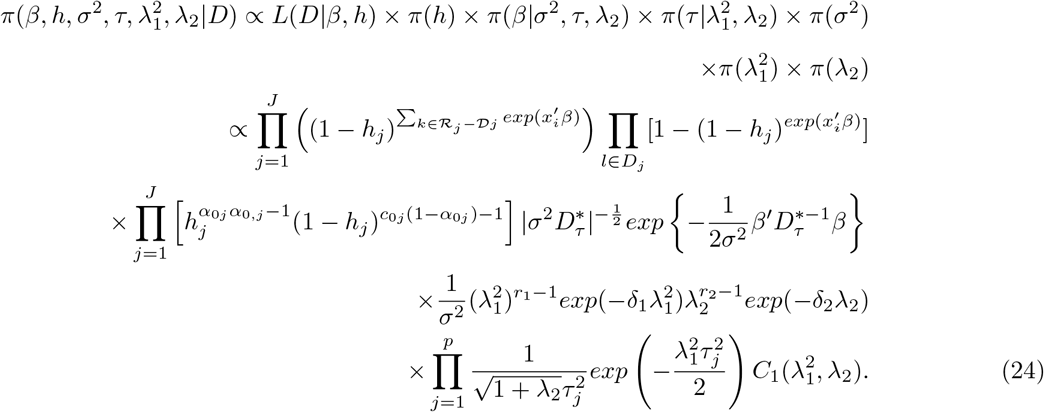

#### 2.3.2. Development PSBC Group Lasso (PSBC-GL-B) joint posterior

The joint posterior distribution for the PSBC Group Lasso (PSBC-GL-B) model, given the cumulative baseline hazard function*H*_0_(*t*), with the beta prior process can be expressed as follows:

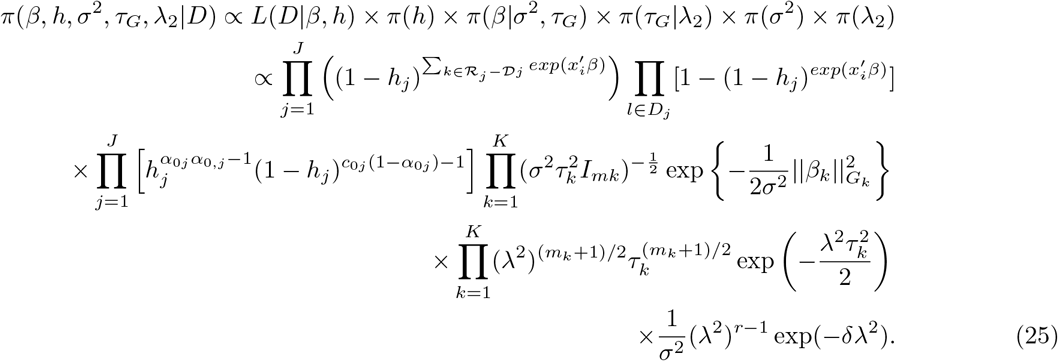

### 2.4. Computational Scheme

This study adopts the Markov Chain Monte Carlo (MCMC) algorithm to fit our proposed models (PSBC-EN-B, Eq. 24, and PSBC-GL-B, Eq. 25) described in the previous section, since the parameters in Equations 24 and 25 cannot be estimated analytically and thus require computational approaches. The computational approach in this study is similar to the method adopted by [40], and the MCMC simulation is used for variable selection, parameter estimation, and prediction in our proposed models (PSBC-EN-B, Eq. 24, and PSBC-GL-B, Eq. 25). We refer the reader to [40] for full details of this computational scheme. However, the steps in Algorithm 1 and 2 depict the step-by-step MCMC algorithms adopted in fitting both the existing and proposed Penalized Semiparametric Bayesian Cox Model (PSBC).

#### Algorithm 1: MCMC algorithm step-by-step procedure for Elastic-Net

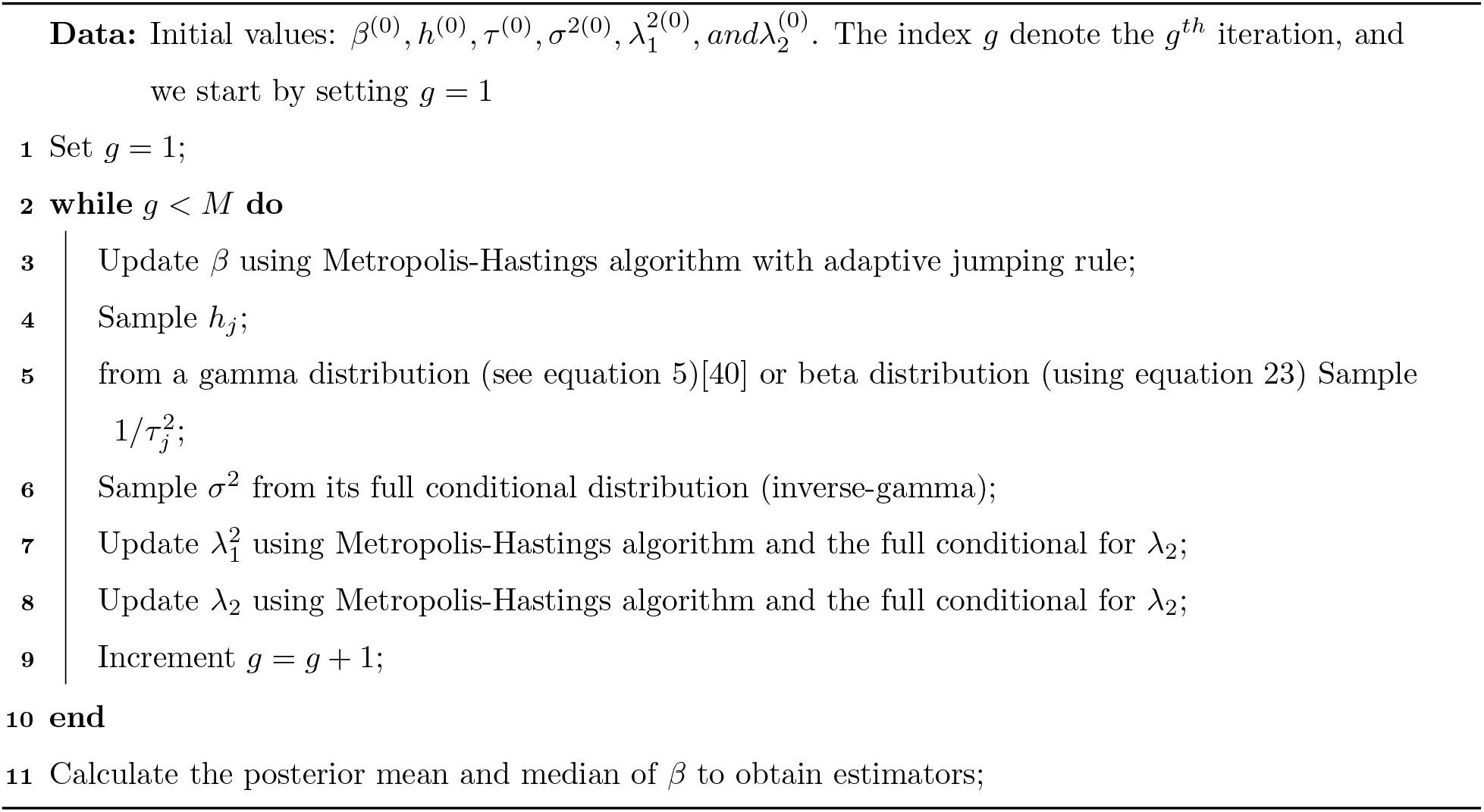

#### Algorithm 2: MCMC algorithm step-by-step procedure for Group Lasso

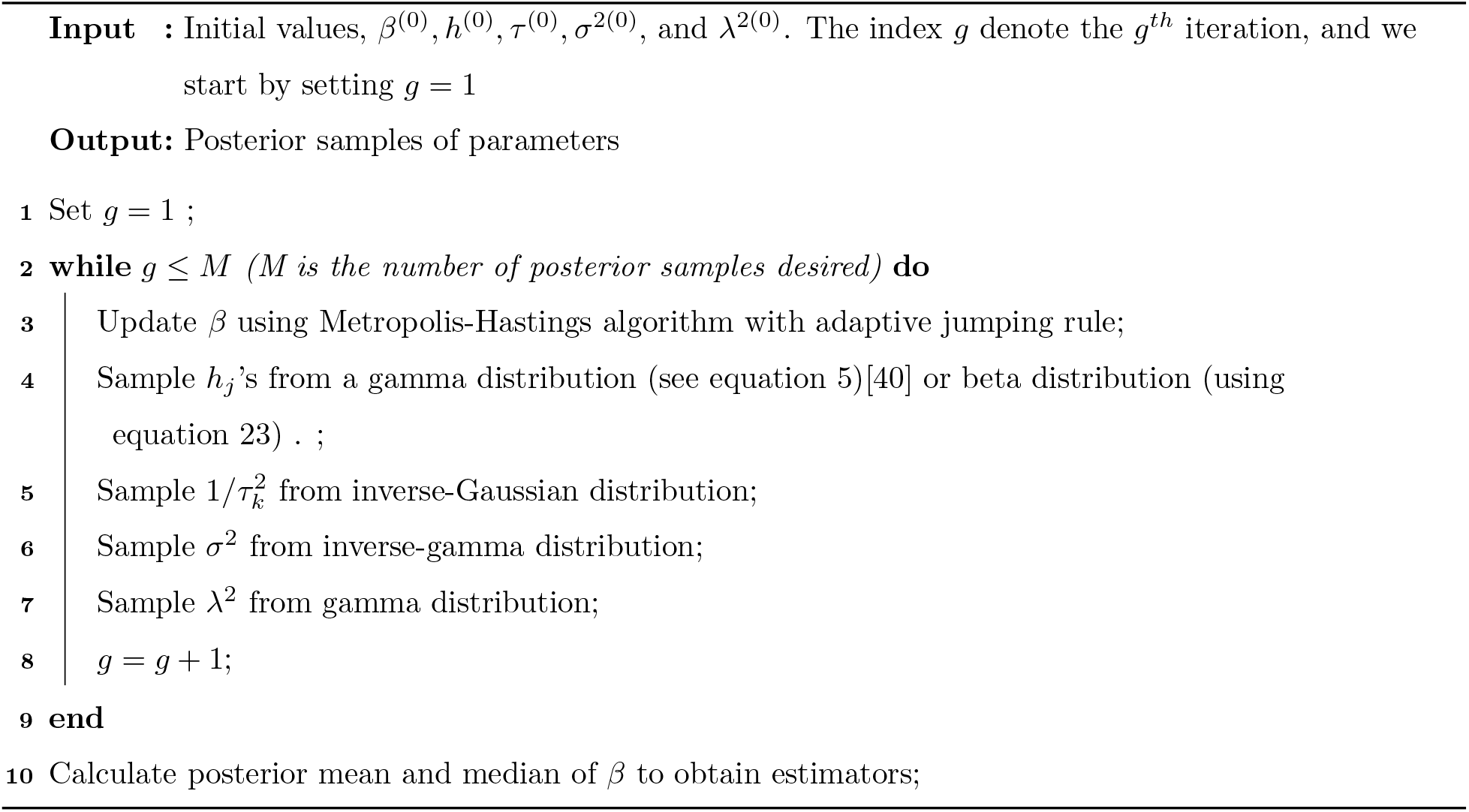

After obtaining the posterior estimates for each method, the sequential BIC were computed using Equation (26) below. The Bayesian information criterion (BIC) is then used as a thresholding method for variable selection in both the proposed and existing models.

### 2.5. Bayesian Information Criterion (BIC)

The Bayesian Information Criterion (BIC) was adopted in the study to choose the variables for the models. After a model’s absolute value estimations of *β*_*j*_ are sorted in descending order, the BIC values are calculated step-by-step by adding significant covariates in equation (26).

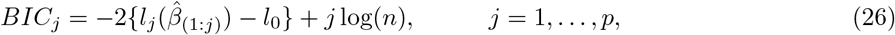

where 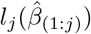 is the maximized log likelihood under a model *M*_*j*_ that includes covariates corresponding to largest *j*|*β*_*j*_|^*′*^s given by 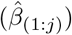, and *l*_0_(0) is the log likelihood under the null model. The least point on the BIC curve refers to the best choice of covariates.

## Results

### 3.1. Simulation Study

In this section, simulation studies was carried out to compare the performance of the methods. A simulation scenario, where a group of relevant covariates are correlated among others while the rest of the covariates are independent of each other were designed.

#### Simulation Procedure

A *ρ* = 0.5 is used to generate the data set. The 10 significant variables, whose coefficients are set equal to 4, are arranged near one another. The study assumed a pairwise correlations with *ρ* = 0.5 only for the 10 important variables. The rest *p −* 10 variables were assumed to be independent of each other. A dataset is generated using *ρ* = 0.5. The 10 significant variables, with coefficients set to 4, are positioned close to each other. The study assumes pairwise correlations of *ρ* = 0.5 for these 10 important variables only, while the remaining *p −* 10 variables are considered independent of each other.

Table 2, report the Bayesian Information Criterion’s (BIC’s) and the average ranks of the BIC value across the PSBC models. The result revealed that the `PSBC-GL-B` with Beta prior gives very impressive results by outperforming the competing models in term of variable selection accuracy. More so, in almost all the cases the PSBC model (`PSBC-GL-B`) consistently gives the least BIC values (Table 2) compared to all other models.

**Table 2:**
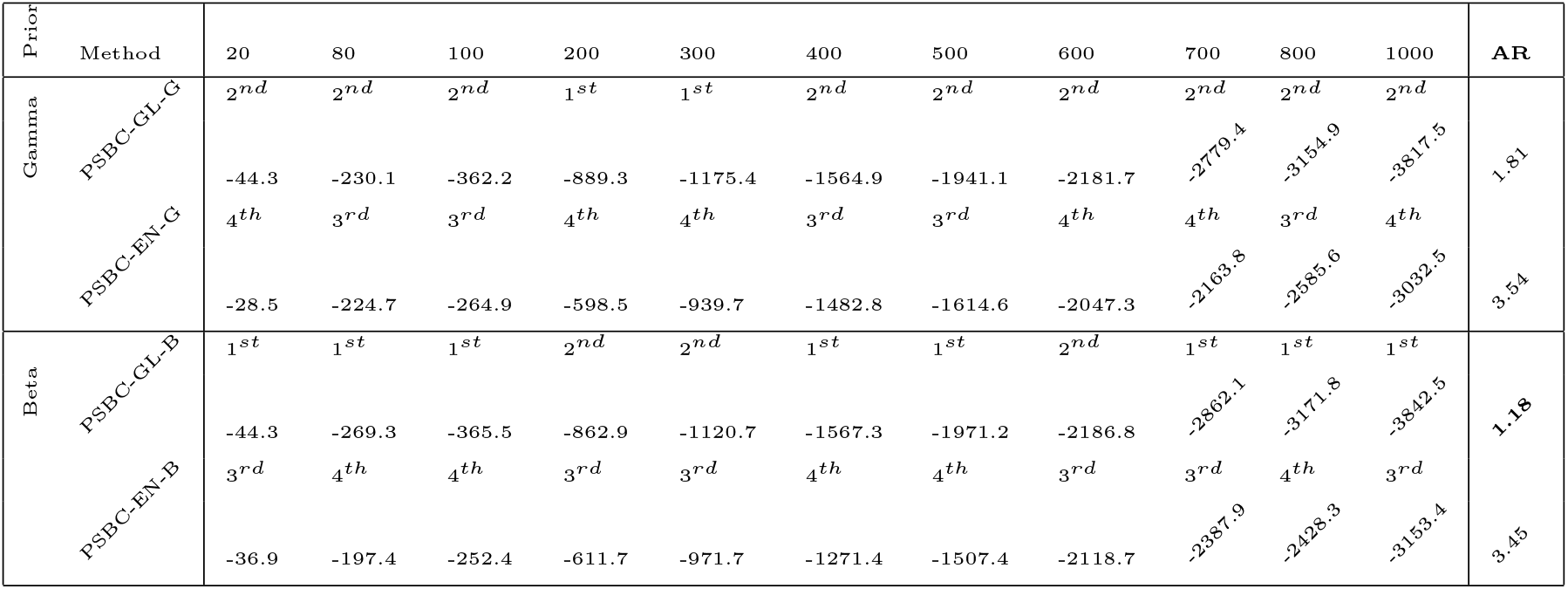
Simulation results based on 100 replications: When 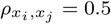, and the Average Rank (AR) of the BIC.

In all, judging by the AR of the method based on the BIC values for each method across the selected number of predictors. We discovered that `PSBC-GL-B` with beta prior was ranked first, `PSBC-GL-G` with *gamma prior* ranked second, followed by `PSBC-EN-B` *beta prior*, and `PSBC-EN-G` *gamma prior*. Table 3 depicts the number predictors selected by the `PSBC` models.

**Table 3:**
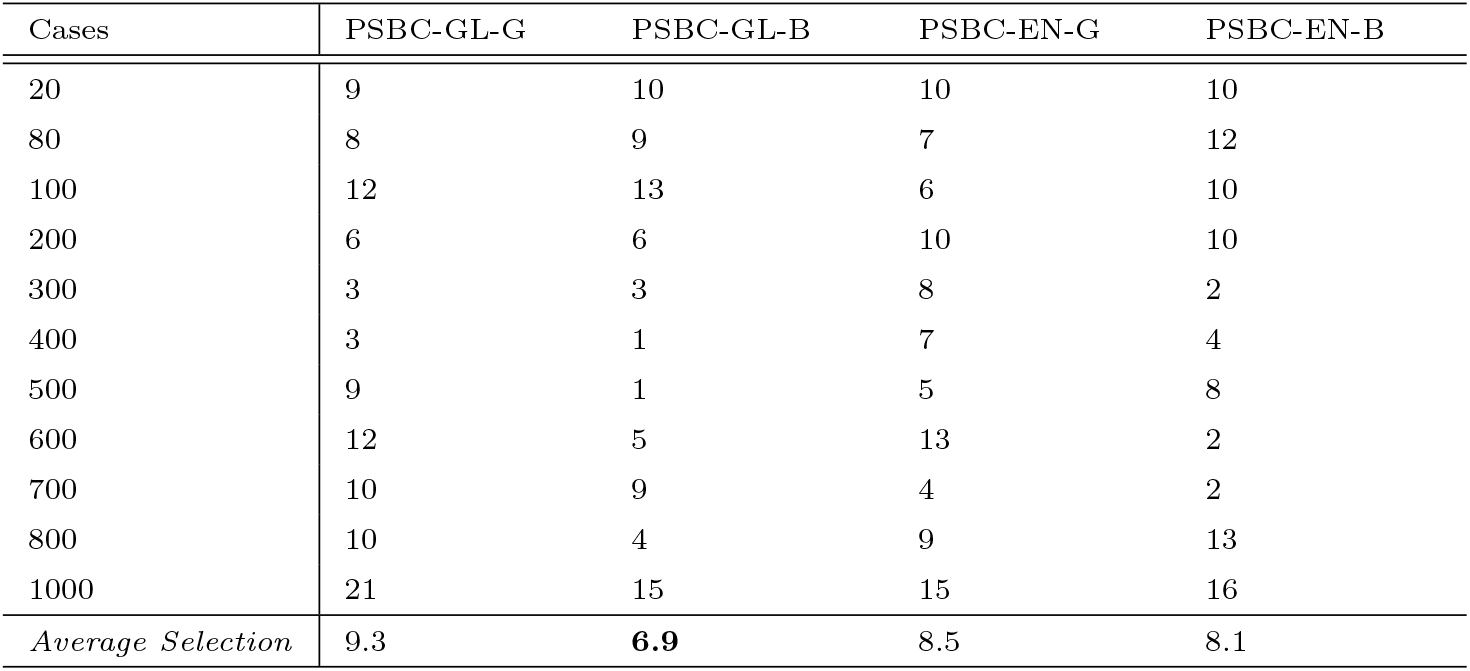
The number of genes selected by PSBC models, using BIC.

| Cases | PSBC-GL-G | PSBC-GL-B | PSBC-EN-G | PSBC-EN-B |
| --- | --- | --- | --- | --- |
| 20 | 9 | 10 | 10 | 10 |
| 80 | 8 | 9 | 7 | 12 |
| 100 | 12 | 13 | 6 | 10 |
| 200 | 6 | 6 | 10 | 10 |
| 300 | 3 | 3 | 8 | 2 |
| 400 | 3 | 1 | 7 | 4 |
| 500 | 9 | 1 | 5 | 8 |
| 600 | 12 | 5 | 13 | 2 |
| 700 | 10 | 9 | 4 | 2 |
| 800 | 10 | 4 | 9 | 13 |
| 1000 | 21 | 15 | 15 | 16 |
| *Average Selection* | 9.3 | **6.9** | 8.5 | 8.1 |

Table 3 revealed that the number of variables or predictors selected by the methods at each instance of predictors in the data simulated. The last row of the table indicate the average number of predictors selected by the methods. The result indicate that modified `PSBC-GL-B` performed reasonably well than the competing methods.

Figure 1 presents the overall performance of the `PSBC` models, using the corresponding BIC values of each model. The plot indicate that the `PSBC-GL-B` outperformed the competing `PSBC` models, followed by `PSBC-GL-G`, and so on.

**Figure 1:**
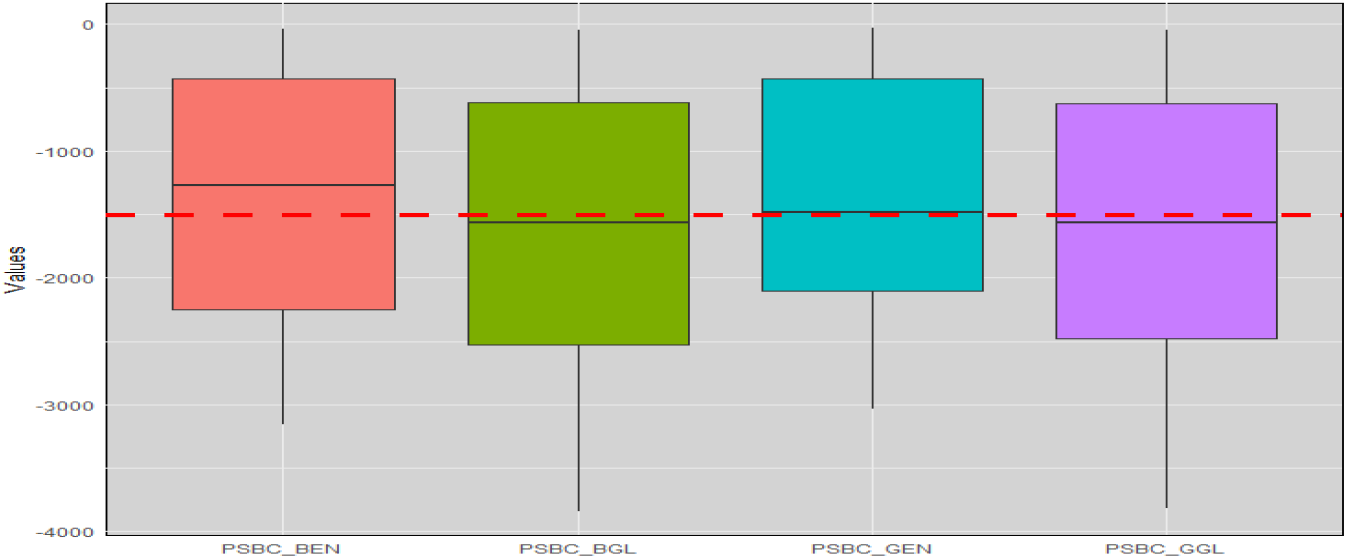
Overall comparison of the PSBC model on the Simulated data set.

### 3.2. Real-Life Studies

This section focused on the practical application of proposed PSBC models along with existing models using three (3) different microarray gene expression data sets. The three data sets are NCI breast cancer data [41] (Data 1), Dutch breast cancer data [42] (Data 2), and Diffuse large-B-cell lymphoma data [43] (Data 3).

A detailed description of the datasets are presented in Table 4. For the first dataset, the number of observations (*n*) are less than predictors (*p < n*) while the remaining two (2) data sets include more predictors than observations (*p > n*).

**Table 4:** The description of three datasets used in this study.

| Data ID | Description | No. of Subjects $n$ | No. of Predictors $p$ |
| --- | --- | --- | --- |
| Data 1. NKI70 | NCI breast cancer data, [41] | 144 | 70 |
| Data 2. DutchBC | Dutch breast cancer data, [42] | 295 | 4919 |
| Data 3. DLBCL | Diffuse large-B-cell lymphoma data patients undergoing R-CHOP treatment, [43] | 181 | 3835 |

The results from Table 4 reveal the dimensions of the three datasets. The first dataset (Data 1: `NKI70`) consists of low-dimensional data since the number of individual genes is fewer than the number of individuals in the dataset. On the other hand, the remaining two datasets are high-dimensional.

### 3.3. Data description using Kaplan-Meier survival curve

To assess the statistical significance of differences between treatment subgroups in the datasets, the study employed the Kaplan-Meier survival curve. This curve was utilized to analyze the duration from the manifestation of cancer symptoms to the incidence of the primary endpoint.

Figure 2 (Left to Right) indicates a significant difference in survival times for the patients at low and high risk groups (`log rank test p=0.018`). The Kaplan-Meier survival probability estimates at 12 months were about 0.76 for the patients at low risk group, and about 0.375 for the patients at high risk.

**Figure 2:**
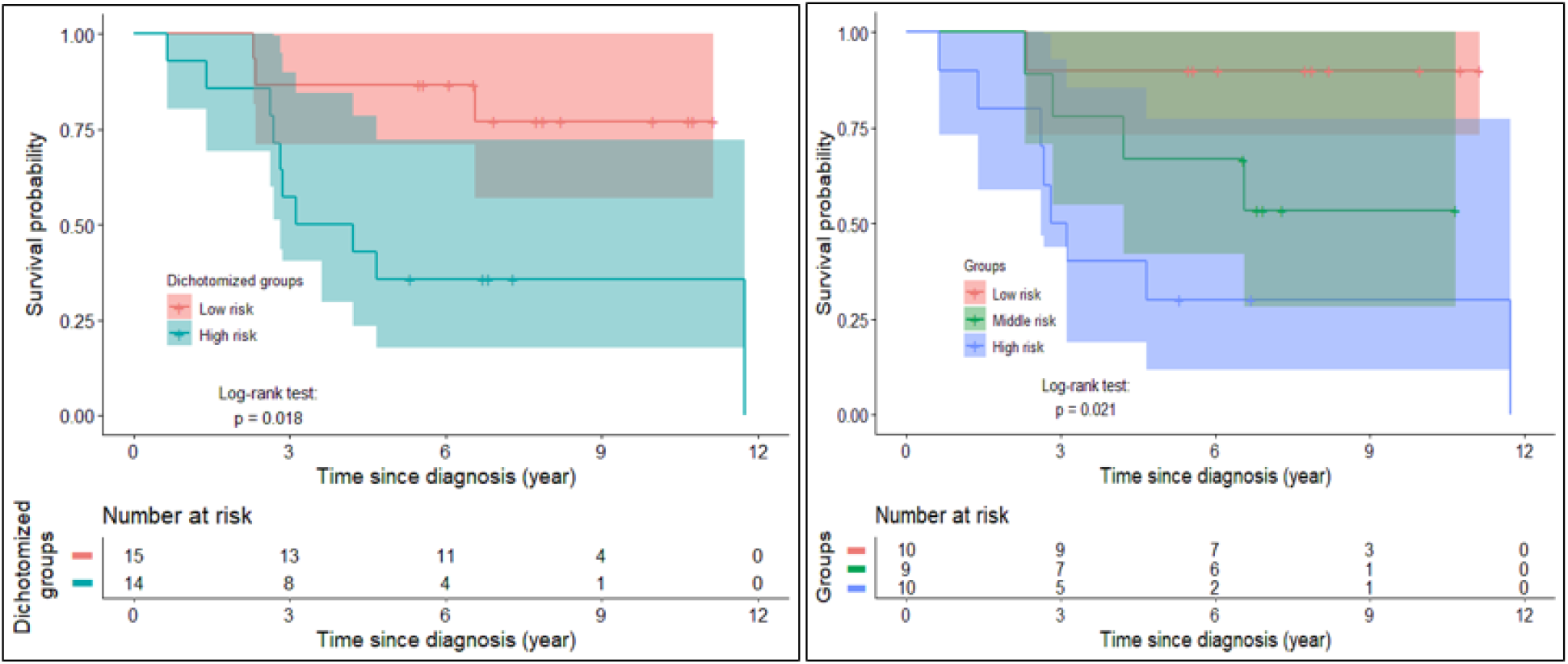
Kaplan-Meier survival curves depict the duration from the onset of breast cancer symptoms to the occurrence of the primary endpoint (death) for different groups. The curves also highlight the ongoing risk of reaching the primary endpoint at various time points, indicating the number of patients still susceptible to the event [Data 1].

As well as in the case where the patients were sub-grouped into three (`Low, Middle`, and `High risk`), a survival difference were noticed between the three (3) subgroups (log rank test p=0.021). More so, the result revealed the the survival probability estimates at 12 months were well above 0.75 for patients at low risk, a little above 0.5 for patient at middle risk, and roughly above 0.25 for patients at high risk.

Figure 3 (Left to Right) indicates a significant difference in survival times for the patients with breast cancer at low and high risk groups (`log rank test p=0.0095`). The Kaplan-Meier survival probability estimates at 12 months were about 0.80 for the patients at low risk group, and about 0.60 for the patients at high risk.

**Figure 3:**
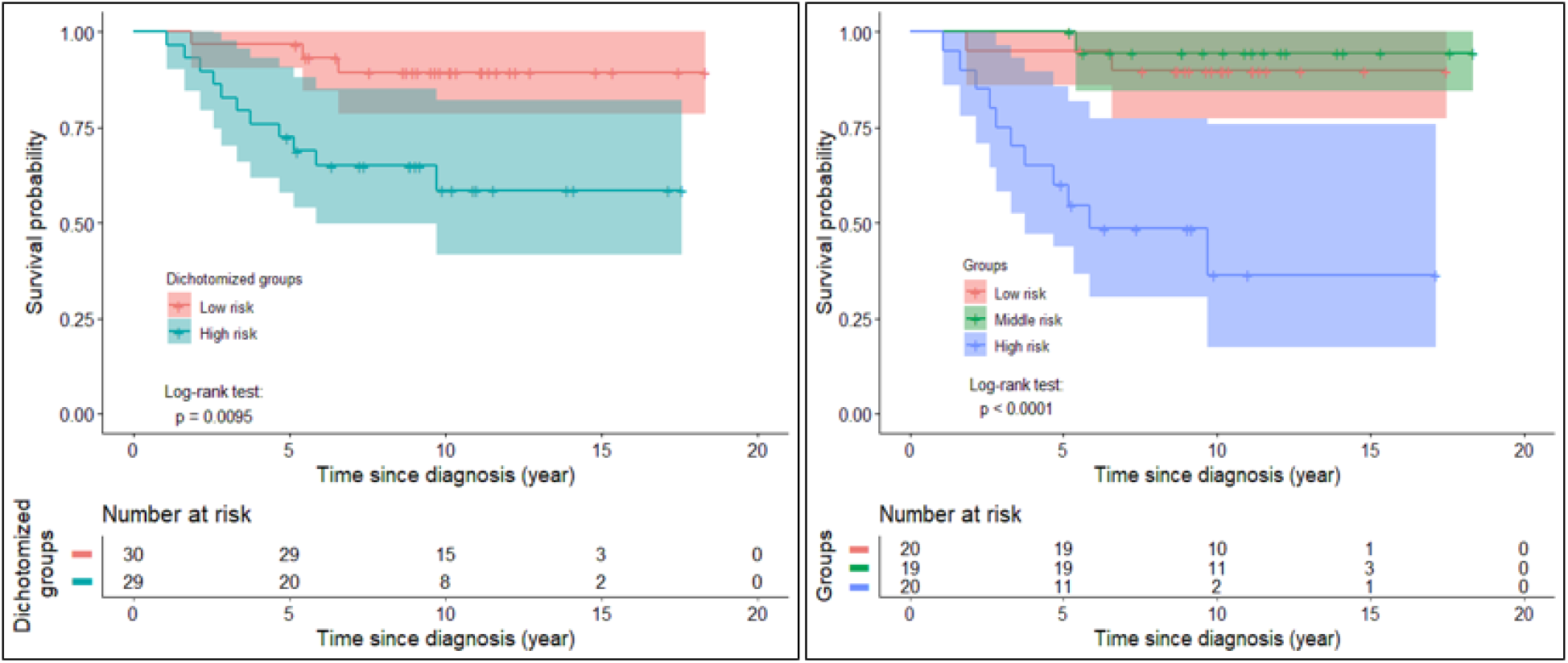
Kaplan-Meier survival curves depict the duration from the onset of breast cancer symptoms to the occurrence of the primary endpoint (death) for different groups. The curves also highlight the ongoing risk of reaching the primary endpoint at various time points, indicating the number of patients still susceptible to the event [Data 2].

Furthermore, in the case where the patients were sub-grouped into three (`Low, Middle`, and `High risk`), a survival difference were noticed between the three (3) subgroups (log rank test p=0.0001). In addition, the survival probability estimates at 12 months were well above 0.90 for patients at low risk, a little above 0.80 for patient at middle risk, and roughly above 0.30 for patients at high risk.

Figure 4 (Left to Right) indicates no significant difference in survival times for the patients with rituxima immunotherapy at low and high risk groups (log rank test p=0.046). The Kaplan-Meier survival probability estimates at 12 months were about 0.80 for the patients at low risk group, and about 0.60 for the patients at high risk.

**Figure 4:**
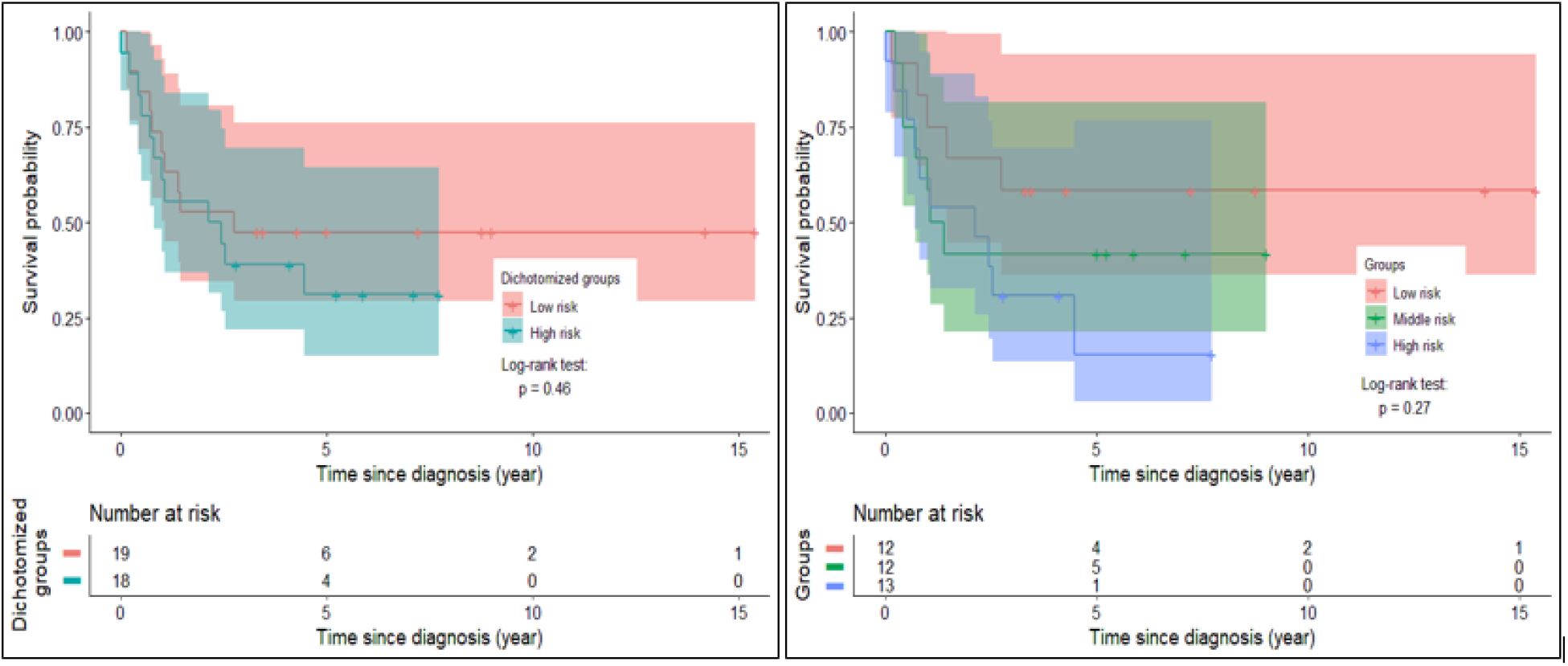
Kaplan-Meier survival curves depict the duration from the onset of rituxima immunotherapy in addition to the chemotherapy symptoms to the occurrence of the primary endpoint (death) for different groups. The curves also highlight the ongoing risk of reaching the primary endpoint at various time points, indicating the number of patients still susceptible to the event [Data 3].

In addition, in the case where the patients were sub-grouped into three (`Low, Middle`, and `High risk`), a survival difference were noticed between the three (3) subgroups (log rank test p=0.27). And the survival probability estimates at 12 months were well above 0.90 for patients at low risk, a little above 0.80 for patient at middle risk, and roughly above 0.30 for patients at high risk.

### 3.4. Comparison of the Methods Performance on the Real-Life data set

This study evaluates the performance of both existing and proposed methods by employing BIC thresholding (Equation 26), which facilitates effective grouped variable selection. Table 5 showcases the comparative results for both the existing and proposed methodologies.

**Table 5:**
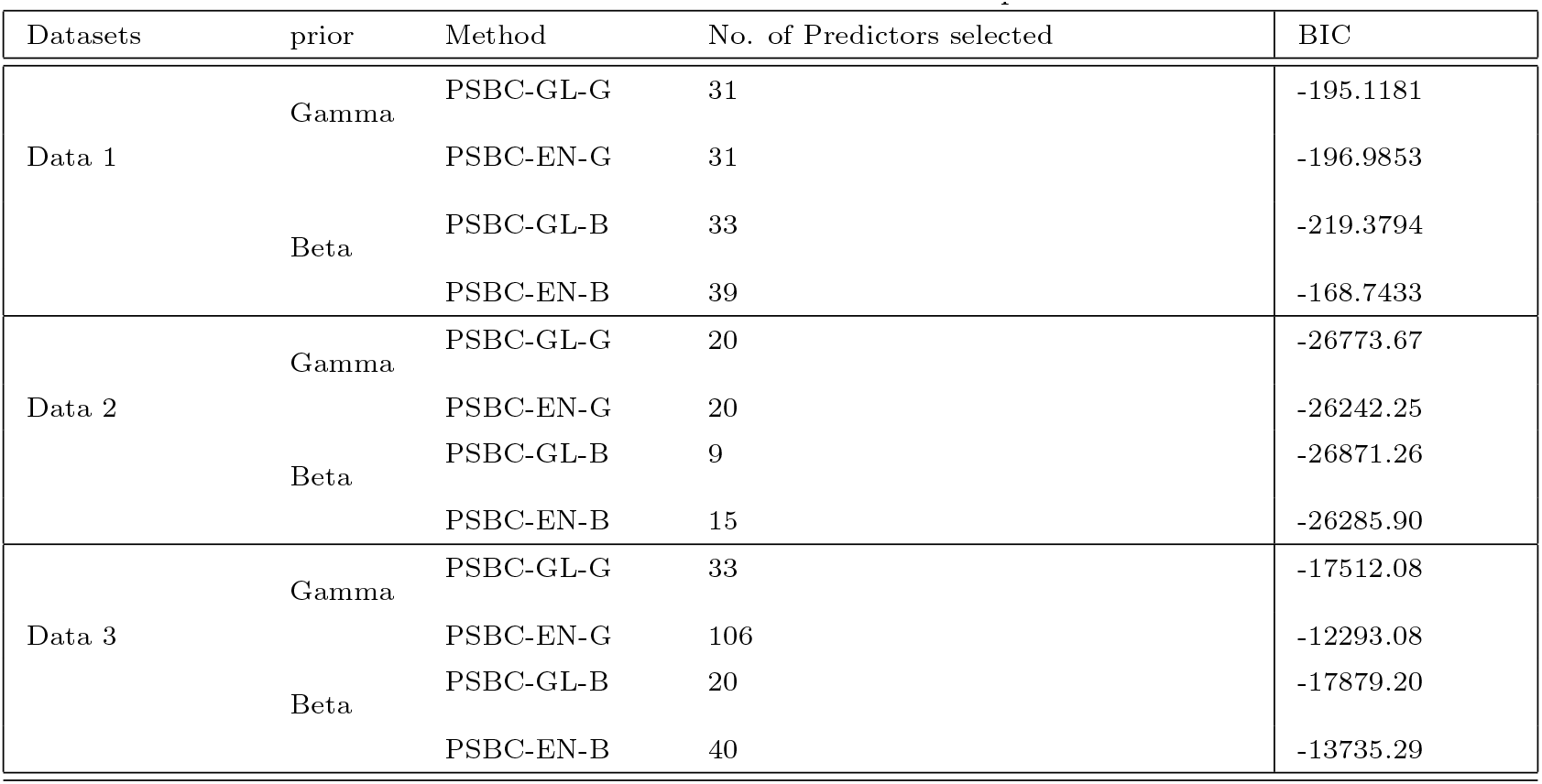
Real-life results based on 100 replications.

Similar to the results from the simulated study, Table 5 indicates that `PSBC-GL-B` demonstrates superior variable selection capability compared to the other three competing methods, as it consistently exhibits the lowest BIC values across the three data sets.

For instance, in the case of `Data 1`, `PSBC-GL-B` achieves the lowest BIC value “**-219.3794**’, followed by `PSBC-EN-G`, with `PSBC-EN-B` performing the least effectively. Similarly, for `Data 2`, `PSBC-GL-B` outperforms the other methods with a BIC of ‘**-26871.26**’, followed by `PSBC-GL-G` ‘**-26773.67**’. The trend continues with `Data 3`, where `PSBC-GL-B` again achieves the lowest BIC value ‘**-17879.20**’.

### 3.5. Posterior Credible Region

In addition, to the results in the previous section on the performance of the methods using the real-life data set. We estimated the credible interval or region for the posterior estimates using both proposed, and the competing methods. However, the Solid dots denote the posterior mode of the coefficients and the lines denote the 95% confidence intervals. The closer the posterior mode to the right side indicates the significant features at a *p <* 0.05. Hence, the plot revealed that few of the features are significantly related to the survival curve.

Figure 5 & 6 depict the posterior credible region for the methods for data 1. From the figures, we noticed that in the case of `PSBC-GL-B` and `PSBC-EN-B` the dots drifted more to the right hand side compare to that of competing methods. This simply implies that the proposed methods are more adequate in more variables in a data set with grouping effect.

**Figure 5:**
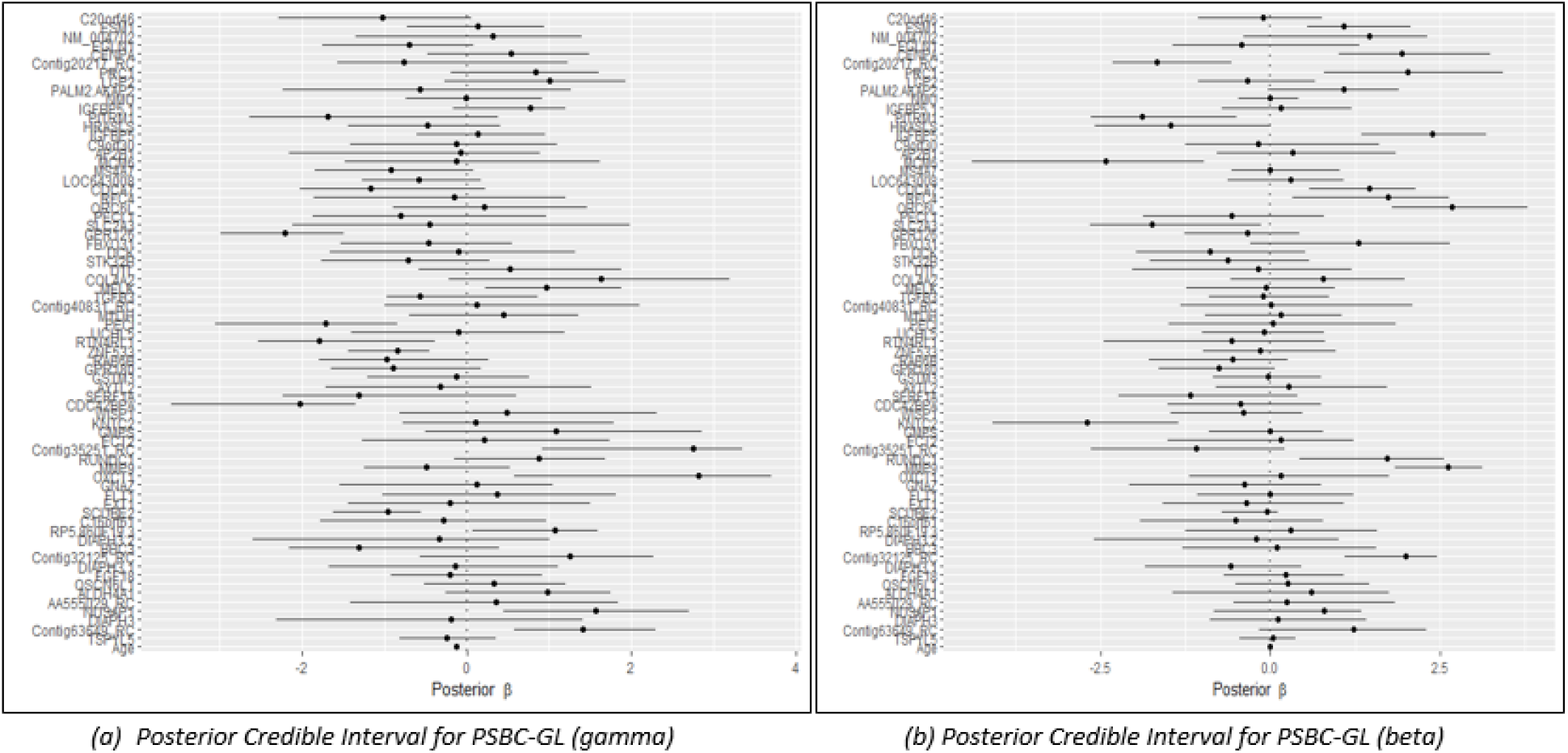
Posterior Credible Region for PSBC-GL: (a) Gamma prior, (b) Beta prior for Data 1

**Figure 6:**
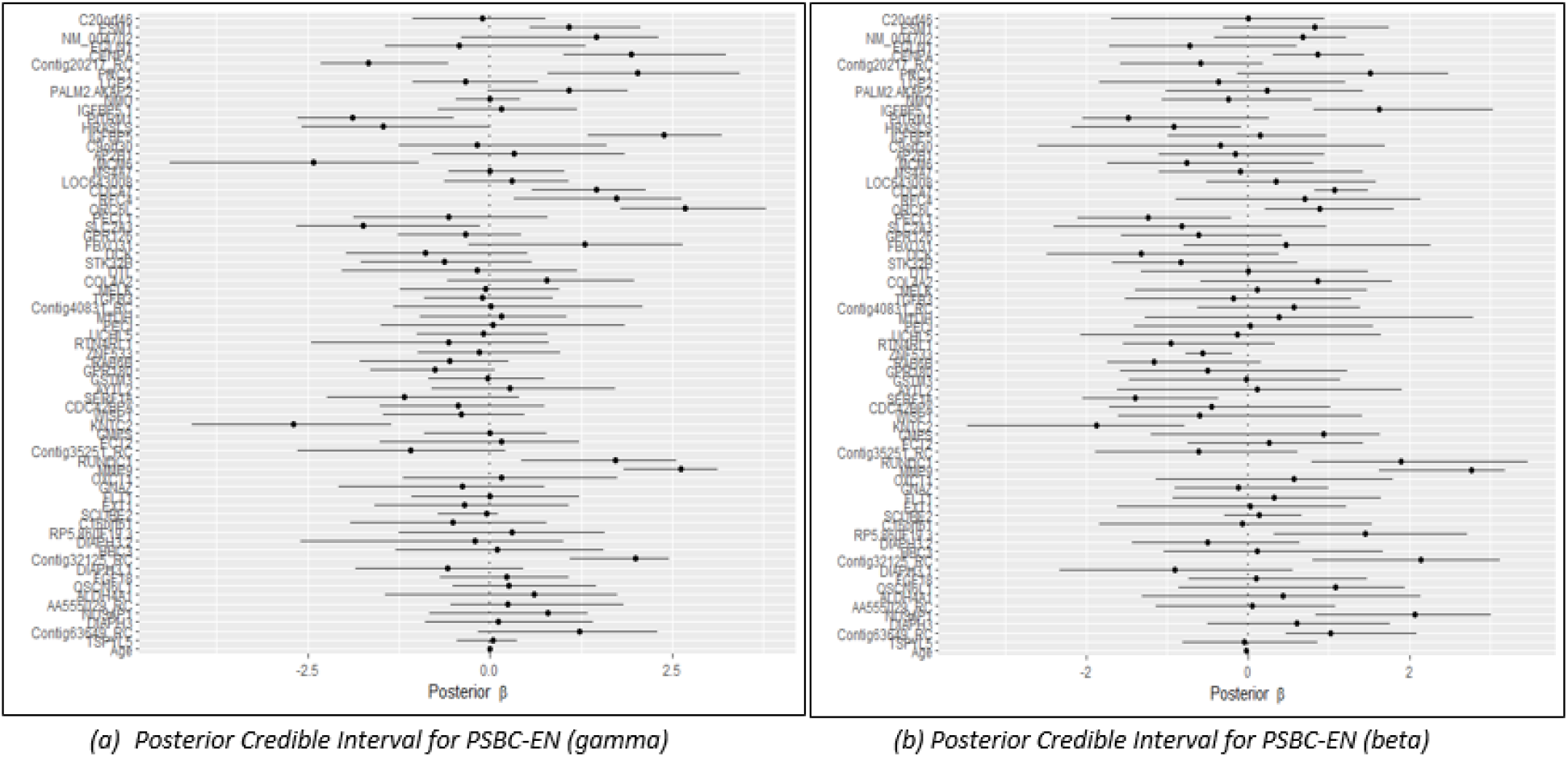
Posterior Credible Region for PSBC-EN: (a) Gamma prior, (b) Beta prior for Data 1

Similar to the findings on Data 1, Figure 7 & 8 represents the posterior credible region for the methods for Data 2, the study revealed that the proposed methods performed better in selecting more variables compared to that of existing methods. Since, the dots of the posterior mode, drifted more to the right hand side of the credible region plots.

**Figure 7:**
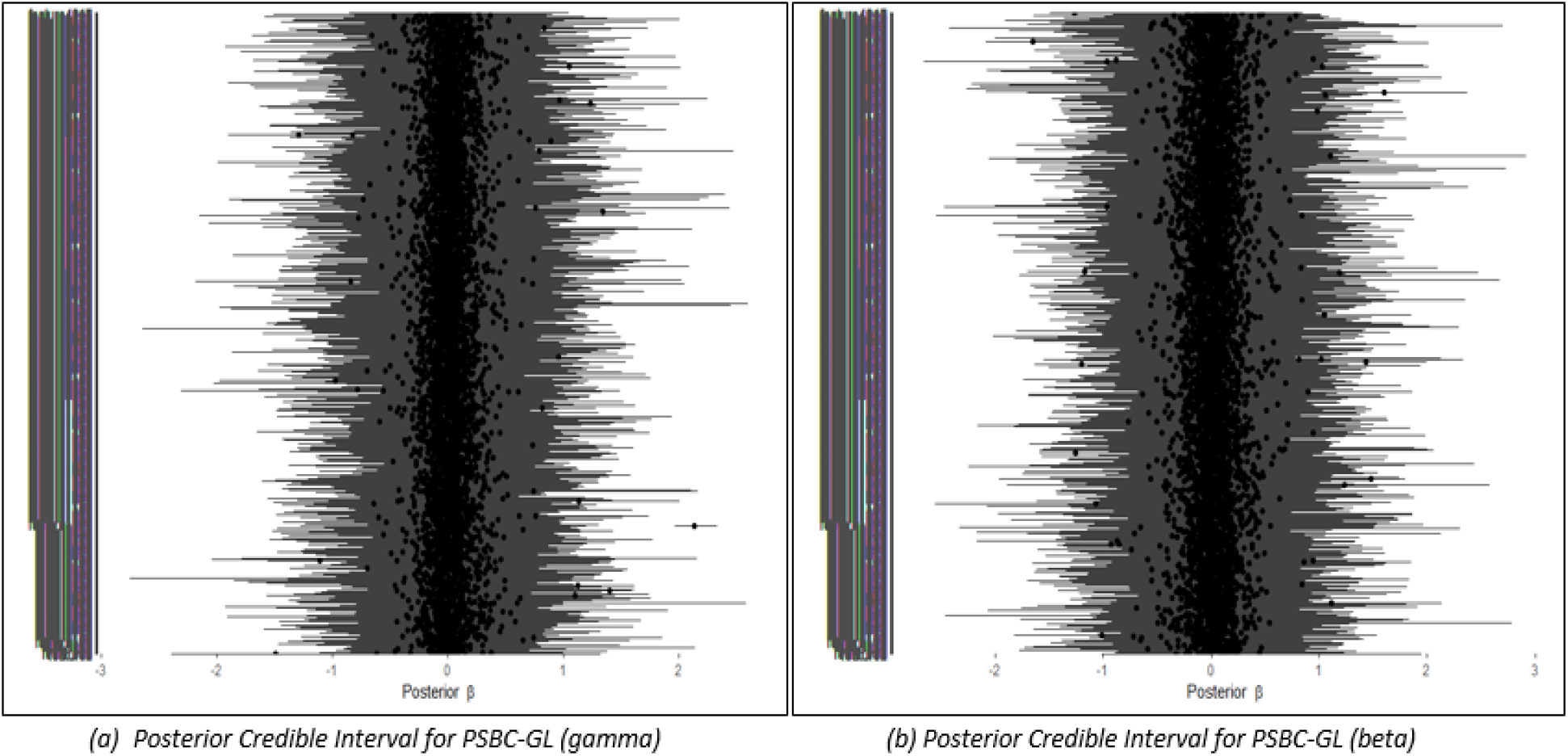
Posterior Credible Region for PSBC-GL: (a) Gamma prior, (b) Beta prior for Data 2

**Figure 8:**
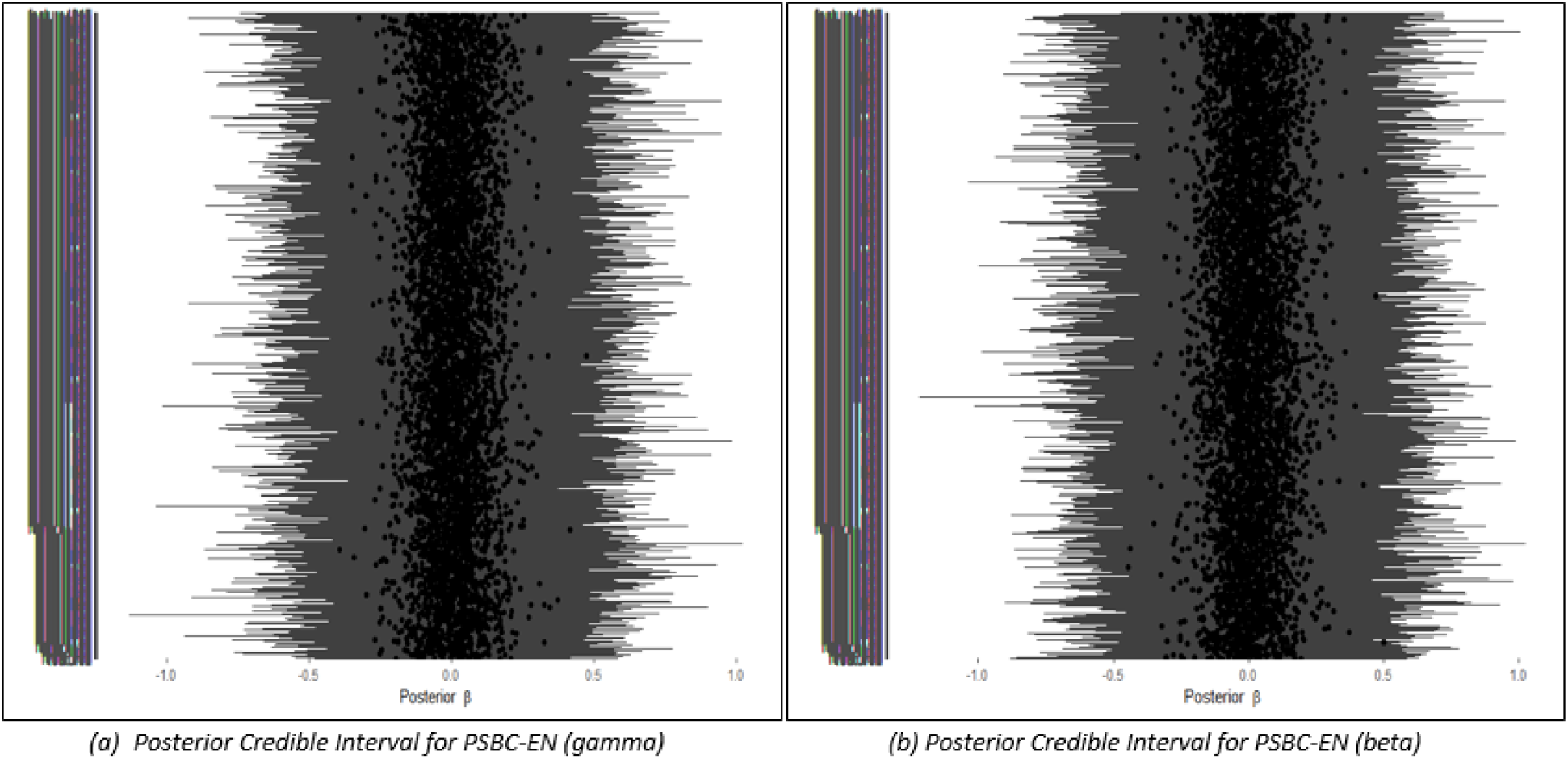
Posterior Credible Region for PSBC-EN: (a) Gamma prior, (b) Beta prior for Data 2

Lastly, in this section, from Figure 9 & 10, it was revealed that few of the features are significantly related to the survival curve for the credible region plots for the proposed methods.

**Figure 9:**
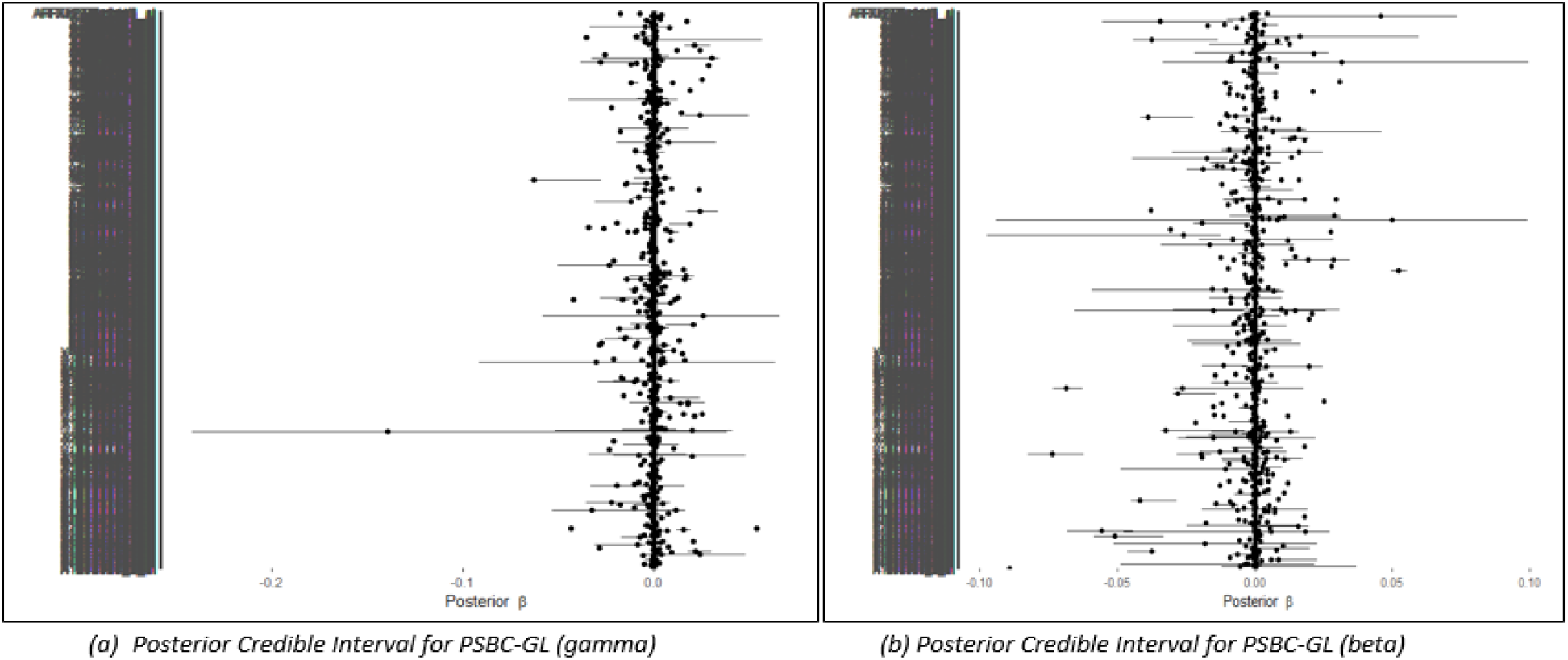
Posterior Credible Region for PSBC-GL: (a) Gamma prior, (b) Beta prior for Data 3

**Figure 10:**
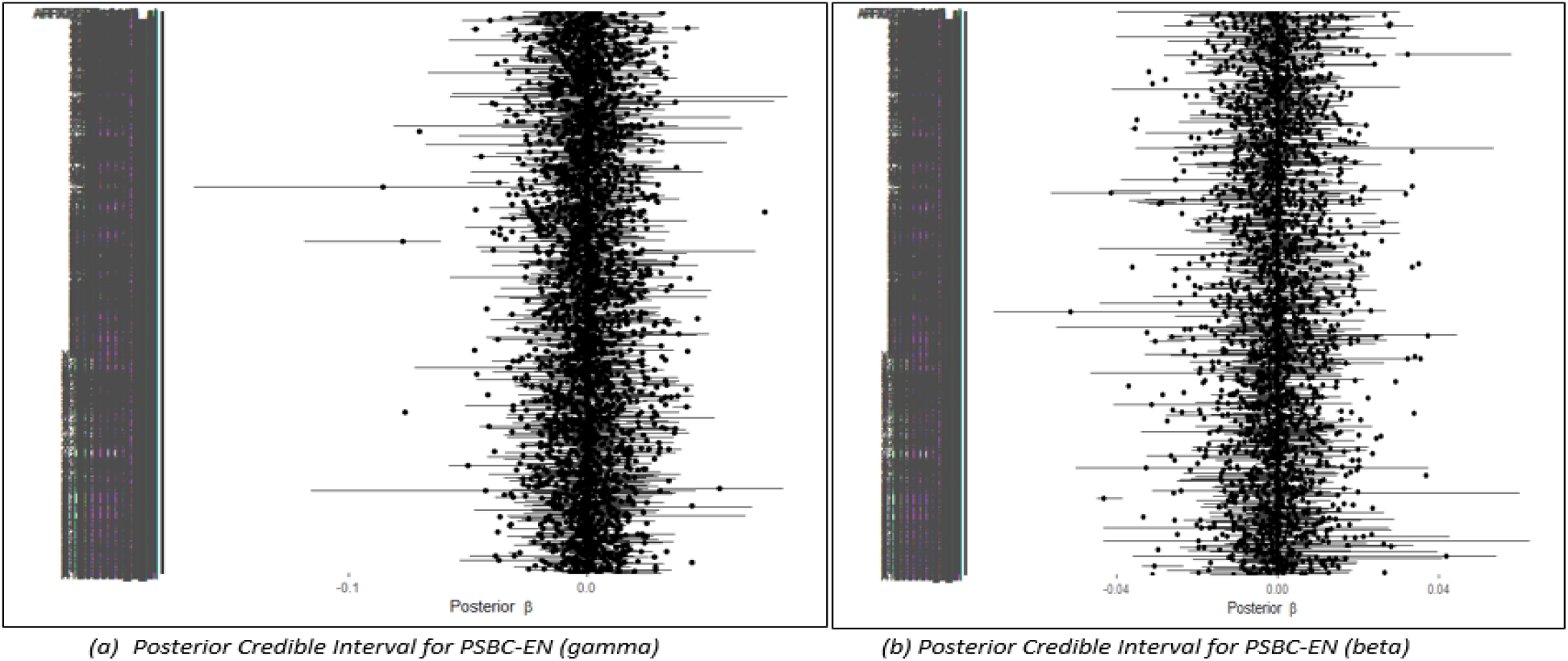
Posterior Credible Region for PSBC-EN: (a) Gamma prior, (b) Beta prior for Data 3

## 4. Conclusions

In this study, we have successfully developed two new penalized Bayesian models for fast variable selection in survival analysis. The new methods (PSBC-GL-B and PSBC-EN-B) are highly efficient in identifying significant covariates, shrinking the coefficients of related (or grouped) variables toward a common value, and, as a result, uncovering any grouping behavior among the covariates, which is a common issue in large scale omics data. Our methodologies implemented a beta process for the cumulative baseline hazard function, contrasting with the gamma process considered in the previous study, thus leading to improvements in the penalized Bayesian model framework.

Through extensive comparative methodologies, the efficacy of our proposed approaches is demonstrated. The results show that the proposed method (PSBC-GL-B) performed better under our simulation settings and real-life datasets in term of variable selection capability and prediction accuracy than the three competing methods, PSBC-GL-G, PSBC-EN-B, and PSBC-EN-G.

## 5. Acknowledgements

Kazeem A. Dauda was supported by the Trond Mohn Foundation (project HyperEvol under grant agreement no. TMS2021TMT09), through the Centre for Antimicrobial Resistance in Western Norway (CAMRIA) (TMS2020TMT11).

